# scPretrain: Multi-task self-supervised learning for cell type classification

**DOI:** 10.1101/2020.11.18.386102

**Authors:** Ruiyi Zhang, Yunan Luo, Jianzhu Ma, Ming Zhang, Sheng Wang

## Abstract

Rapidly generated scRNA-seq datasets enable us to understand cellular differences and the function of each individual cell at single-cell resolution. Cell type classification, which aims at characterizing and labeling groups of cells according to their gene expression, is one of the most important steps for single-cell analysis. To facilitate the manual curation process, supervised learning methods have been used to automatically classify cells. Most of the existing supervised learning approaches only utilize annotated cells in the training step while ignoring the more abundant unannotated cells. In this paper, we proposed scPretrain, a multi-task self-supervised learning approach that jointly considers annotated and unannotated cells for cell type classification. scPretrain consists of a pre-training step and a fine-tuning step. In the pre-training step, scPretrain uses a multi-task learning framework to train a feature extraction encoder based on each dataset’s pseudo-labels, where only unannotated cells are used. In the fine-tuning step, scPretrain fine-tunes this feature extraction encoder using the limited annotated cells in a new dataset. We evaluated scPretrain on 60 diverse datasets from different technologies, species and organs, and obtained a significant improvement on both cell type classification and cell clustering. Moreover, the representations obtained by scPretrain in the pre-training step also enhanced the performance of conventional classifiers such as random forest, logistic regression and support vector machines. scPretrain is able to effectively utilize the massive amount of unlabelled data and be applied to annotating increasingly generated scRNA-seq datasets.

**Availability:** https://github.com/ruiyi-zhang/scPretrain\

## 1 INTRODUCTION

Recent breakthroughs in single cell sequencing technology have generated a large number of large-scale scRNA-seq datasets[1–10], which hold the promise of understanding cellular differences and the function of each individual cell at single-cell resolution. One of the most important steps in single-cell analysis is to classify each cell into its cell type based on the gene expression profile. Manually labelling cell types cannot scale to massively expanding single cell datasets and also requires domain knowledge for specific tissues and organs. For example, one of the existing largest single cell datasets was annotated by a group of 39 domain experts[11]. To accelerate the labor-intensive manual curation, supervised learning approaches have been used to automatically classify cells using existing annotated cells[12–20]. Despite the encouraging performance of these approaches, they often ignore unannotated cells, which are more abundant than annotated cells and contain rich information of gene activity. Moreover, limiting the analysis to annotated cells can also lead to overfitting and being vulnerable to batch effects[21–24], hindering progress towards comprehensive cell type annotation and cellular diversity understanding. Intuitively, jointly leveraging both unannotated and annotated cells might collectively address these problems and thus advance cell type classification.

Self-supervised learning (SSL) approaches have obtained the state-of-the-art results in various machine learning fields like natural language processing[25], video analysis[26] and computer vision[27]. SSL often consists of a pre-training step and a fine-tuning step. In the pre-training step, SSL extracts high-quality feature representations using only the unannotated data. In the fine-tuning step, these representations are used to train a supervised model for the specific task of interests. For example, BERT used masked language model and next sentence prediction to learn a feature encoder in the pre-training step[25]. After that, task-specific inputs and outputs were used to fine-tune the feature encoder in the fine-tuning step. These two steps collectively leverage the abundant unannotated data and expensive annotated data to achieve better classification performance. Unlike semi-supervised learning approaches[28–32], SSL does not leverage annotated data in the pre-training step, thus generating more robust features for downstream applications.

Despite the success of SSL in other machine learning areas, pre-training on gene expression data remains challenging due to at least two reasons. First, different from word sequences[25], graphs[33] or images[34] that have explicit topological structure, gene expression feature matrix does not have ordered structure. This ordered structure, however, is the key element in SSL methods such as BERT which masks and predicts a word using nearby context. Second, single cell datasets present substantial batch effects due to different technologies, platforms, animals, organs and species[21–24]. Such batch effects could prevent the feature extraction encoder from learning underlying gene expression patterns. These batch effects could be further amplified when we do pre-training on a large number of diverse datasets. Collectively, pre-training on unordered and diverse gene expression datasets remains challenging.

In this paper, we proposed a novel multi-task self-supervised learning approach scPretrain for cell type classification. The key idea of scPretrain is to use a multi-task learning framework which views training on each dataset as a single pre-training task, thus preventing the noise from batch effects. Moreover, since gene expression matrix does not have any ordered structure, scPretrain first uses K-means to cluster unannotated cells from each dataset and then trains a feature extraction encoder by using cluster labels as pseudo-labels. The clustering and classification steps are performed iteratively to enhance the feature extraction encoder. We extensively tested scPretrain on 60 datasets and obtained an average improvement of 7.3% on AUPRC and 5.3% on AUROC compared against neural networks without pre-training. We further showed that the representations created by our model can also improve the performance of other off-the-shelf classifiers. scPretrain is able to effectively utilize the massive amount of unlabelled scRNA-seq datasets to advance cell type classification and single cell analysis.

## 2 METHODS

The goal of the pre-training step is to learn a feature extraction encoder, which would be used in the fine-tuning step to classify new cells. We first used K-means to cluster cells in each single cell dataset, resulting in a pseudo-label for each cell. After that, we exploited a multi-task learning framework to train a feature extraction encoder shared by different datasets, in order to alleviate batch effects. We then iteratively leveraged new cell representations to refine pseudo-labels, which were further used to retrain the multi-task learning framework. In the fine-tuning step, the pre-trained encoder can be used on downstream tasks like classification and clustering.

**Fig. 1.**
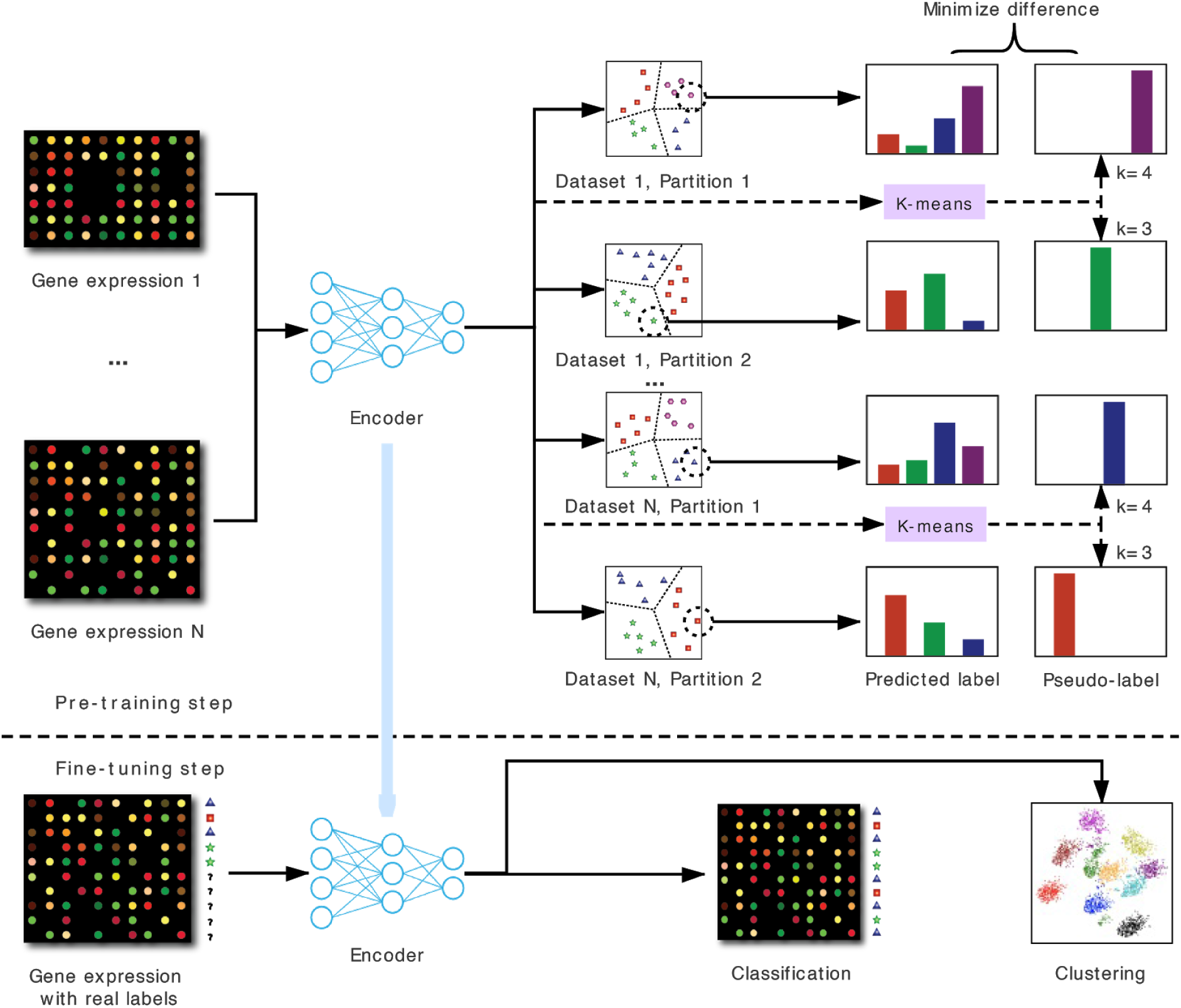
Flowchart for scPretrain. In the pre-training step, scPretrain assigns a pseudo-label to each cell using K-means. These pseudo-labels are used to train a feature extraction encoder, which is shared by different datasets and different partitions in a multi-task learning framework. In the fine-tuning step, this encoder is used to embed cells in a new dataset to low-dimensional representations, which are further used in downstream tasks such as cell clustering and cell type classification.

### 2.1 Problem definition

scPretrain consists of a pre-training step and a fine-tuning step. In the pre-training step, the input is a collection of unlabelled datasets, 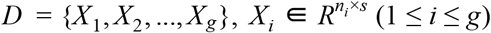. Here, *X*_*i*_ is a gene expression matrix of *n*_*i*_ cells and *s* genes. The goal in the pre-training step is to learn a feature extraction encoder *E*_θ_ (*x*) that maps a high-dimensional gene expression vector *x* ∈ *R*^*s*^ of a cell to a low-dimensional vector *h* ∈ *R*^*r*^, where *r* is the dimension of the cell representation. This feature extraction encoder is shared across all unannotated datasets in the pre-training step. In the fine-tuning step, the input is an annotated dataset *Z* ∈ *R*^*m*×*s*^, *Y* ∈ *R*^*m* ×*k*^, where *Z* is the gene expression matrix of *m* cells and *s* genes. *Y* is the binarized cell type label matrix and *k* is the number of cell types. *Y*_*ij*_ = 1 if the *i*-th cell is cell type *j*, otherwise *Y*_*ij*_ = 0. The goal at the fine-tuning step is to train a cell type classifier using both the feature extraction encoder *E*_θ_ from the pre-training step and annotations from *Y*. This classifier will be evaluated on another test dataset *Q*, where the goal is to predict the cell type of cells in *Q*. The feature extraction encoder *E*_θ_ is shared across all dataset in the pre-training and fine-tuning steps, thus forcing it to encode the common gene expression patterns rather than batch effects in individual datasets. As a consequence, scPretrain is scalable to the large number of diverse unannotated datasets and can be rapidly used to annotate new datasets.

### 2.2 Pre-training using pseudo-labels

Given a gene expression matrix *X* ∈ *R*^*n*×*s*^ of *n* cells and *s* genes, our goal is to learn a feature extraction encoder *E*_θ_ (*x*), which is expected to perform well on other datasets in various downstream tasks. Let *Y′* ∈ *R*^*n*×*k*′^ be the labels of the training set. Let *F*_ω_(*h*) = *F*_1_(*sigmoid*(*h*)) be the classifier, where 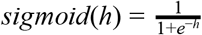, *h* is the output of the encoder *E*_θ_, and *F*_1_ is a fully-connected neural network with output dimension *k*′. Let *x* be an input gene expression vector, *t* = *F*_ω_(*E*_θ_(*x*)) be the corresponding *k*′ dimension output vector and *y* be the binary label vector of *x*, *v*_*i*_ is the predicted score that *x* belongs to class *i*:

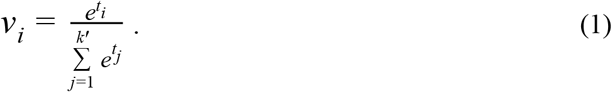

We use the cross entropy loss 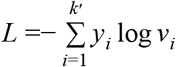 as the loss function. Therefore, we can optimize the following function to get *E*_θ_ (*x*):

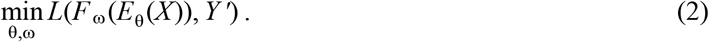

However, optimizing the above loss function is unfeasible since we do not have the ground truth labels *Y*′ for cells in *X*. We propose to use self-supervised learning to train the feature extraction encoder *E*_θ_ (*x*). In particular, we use pseudo-labels as an alternative to ground truth labels. Pseudo-label methods first create pseudo-labels directly from raw input data and then use these pseudo-labels to train the feature extraction encoder[27,35]. When real labels are available, these real labels are then used to fine-tune the feature extraction encoder. Pseudo-label methods assume that samples are from different underlying subgroups and each subgroup corresponds to a well-defined class. Pseudo-labels approximate these underlying groups and thus guide us to train a good feature extraction encoder. Notably, this process of utilizing pseudo-labels mimics the process of manual cell type curation, where cells are first clustered into different groups according to the gene expression profile, then each group is assigned with one cell type by experts [11,36]. We hope to learn a good feature extraction encoder by finding the underlying patterns that lead to different cell subgroups. We select K-means as our base clustering algorithm. In practice, we found that using other clustering methods (e.g. hierarchical clustering[35]) yielded similar performance.

The pre-training step has two phases. In the first phase, formally, we define a centroid matrix *C* with size 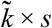, where *s* is the input dimension, and the pseudo-label matrix 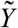 with size 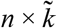, which can be regarded as the cluster assignment of each cell. We then do clustering by optimizing the following loss function simultaneously on *C* and 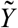:

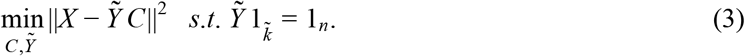

We will use 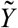, but we do not use *C* directly. The K-means algorithm will create 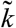 centroids optimizing formula (3). Recent unsupervised single cell clustering methods show good alignment between ground truth cell type labels and cell clusters[36]. Likewise, each centroid obtained by K-means might reflect the underlying cell type and can thus be used as an alternative to ground truth cell types to train a good feature extraction encoder. We then learn the feature extraction encoder *E*_θ_ using pseudo-labels 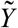 by optimizing the following function:

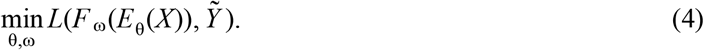

In the first phase mentioned above, we obtain pseudo-labels by clustering on the origin gene expression profile. After having the first batch of pseudo-labels, we can train the feature extraction encoder that maps the high-dimensional gene expression profile into low-dimensional cell representation. Consequently, in the second phase, we obtain pseudo-labels by clustering on the resulting cell representations. The first phase is only performed once, whereas we iteratively perform clustering and classification in the second phase until convergence. This iterative process further enhances the feature extraction encoder by boosting the discriminative power of the extracted features. We still use K-means to create pseudo-labels on encoder output features:

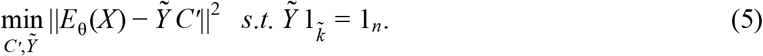

Here, θ is fixed during this clustering procedure and the size of centroid matrix *C*′ becomes 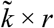, where *r* is the encoder output dimension, and 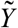 is still the pseudo-label matrix. Although clustering on encoder outputs does not add any additional information to the model, the following classification steps optimize encoder parameters and force the representation of each cell to be as close as possible to the centroid of its cluster. The classification procedure is still the same as formula (4).

### 2.3 Multi-task pre-training

Batch effects make gene expression profiles vary largely in distribution. We use a multi-task learning framework to address this problem. Given *g* datasets *D* = {*X*_1_, *X*_2_, ..., *X*_*g*_} and their pseudo-label set 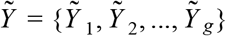, we minimize the pseudo-label based cell type classification loss on *g* datasets simultaneously. Specifically, each dataset uses the same encoder *E*_θ_, but they have *g* different classifiers *F* = {*F*_1_, *F*_2_, ..., *F*_*g*_}. We use ω = {ω_1_, ω_2_...ω_*g*_} to denote the parameters in *g* classifiers. We estimate the parameters of the encoder θ by optimizing the following problem:

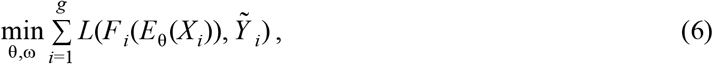

where *L* is the cross entropy loss function mentioned above. This multi-task learning framework forces the feature extraction encoder *E*_θ_ to map gene expression profiles to a robust representation which is less affected by batch effects from different datasets.

We implemented a two phase pre-training process similar to section 2.2 to create pseudo-labels and trained the encoder on them. Firstly we create the centroid matrix set *C* = {*C*_1_, *C*_2_, ..., *C*_*g*_} and label assignment 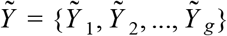 for each dataset by optimizing the following function:

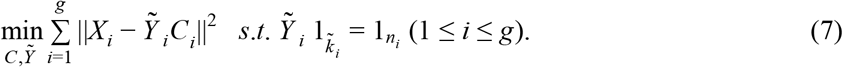

Here each matrix *C*_*i*_ in *C* has the shape 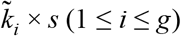, where 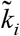 is the number of pseudo-label types set by the dataset quantity mentioned in section 2.2. And each dataset *X*_*i*_ in *D* has the shape *n*_*i*_ × *s* (1 ≤ *i* ≤ *g*). After we have the pseudo-labels created by formula (7), we feed these data to formula (6) and optimize that function in order to estimate the encoder *E*_θ_. Then we start our second pre-training step using this encoder, which updates the pseudo-label creating function to:

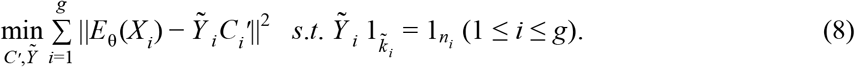

At this time, each centroid matrix *C*_*i*_′ in *C*′ = {*C*_1_′, *C*_2_′...*C*_*g*_′} has the shape 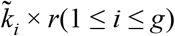. Through optimizing formula (8) and (6) iteratively, we take turns to update the parameters θ and 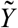 on all the datasets to finally obtain the feature extraction encoder.

### 2.4 Determining the number of clusters in each task

A key hyperparameter for K-means is the number of clusters 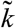, which is empirically difficult to predefine. We thus use different 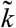 to cluster each dataset and treat each partition as a single task in our multi-task learning framework. Consequently, our multi-task learning framework would have *g* × *p* tasks, where *p* is the number of partitions. The intuition behind considering different 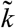 here is to force the encoder *E*_θ_ to become insensitive to the number of clusters in a new dataset.

Formally, the centroid matrix sets in the two phases now become *C* = {*C*_11_, *C*_12_, ..., *C*_*gp*_} and *C*′ = {*C*_11_′, *C*_12_′, ..., *C*_*gp*_′}. The pseudo-label matrix set becomes 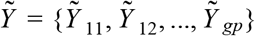. As a result, the new clustering loss function in each of the two phases become:

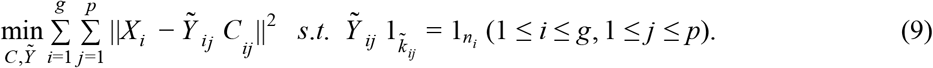

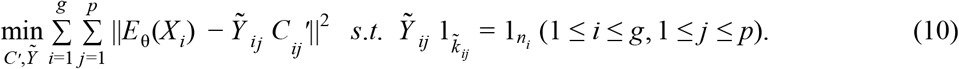

The classification function is similar to formula (6) by predicting 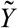. A recent study proposed to find a more robust number of clusters by using a validation set[35]. In contrast to this approach, we take the advantage of the multi-task learning framework to thoroughly examine different 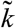. In practice, we set the average data point number in each cluster to be 50, 100 and 200, then calculated the related three cluster numbers of each dataset as the candidates for 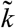.

### 2.5 Fine-tuning for cell type classification

In the fine-tuning step, we fine-tuned the pre-trained feature extraction encoder on a new dataset with real labels. We used a deeper classifier *F*_ϕ_(*h*) = *F*_2_(*sigmoid*(*F*_3_(*sigmoid*(*h*)))), where *F*_2_, *F*_3_ are fully-connected neural networks. In contrast, we use a less expressive neural network in the pre-training step in order to avoid overfitting by using pseudo-labels. We optimized the following function to estimate the parameters using these real labels:

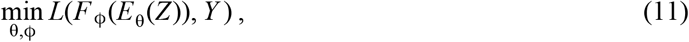

where *Y* is the real label matrix, instead of the pseudo-labels generated. If we want to use other off-the-shelf classifiers instead of neural networks, we will use the cell representation *h* = *E*_θ_(*x*) instead of *x* as the input for other classifiers such as logistic regression and random forest.

## 3 EXPERIMENTAL RESULTS

### 3.1 Datasets

We obtained 60 scRNA-seq datasets from Cell BLAST[37]. The organ and platform selection was not limited to any certain type, while we only chose mouse and human datasets in our experiments. These 60 datasets span over 29 organs and 10 platforms, reflecting substantial batch effects among them. The number of genes in each dataset varied from 870 cells (Quake Smart-seq2 Diaphragm) to 160,796 cells (Zeisel 2018) with an average number of 15,209 cells. The number of cell types in each dataset was between 2 and 81, with an average of 10 cell types. To map genes across species, we first unioned all the genes in datasets of each species, then found the intersection of mouse genes and human genes, which contained 17,298 genes. We performed pre-training on the expression vector of these 17,298 genes. We used zero imputation for genes that were missing in a specific dataset. In the fine-tuning step, we adjusted the weight matrix based on pre-training genes, and used random initialization to the parameters of those genes not included in these 17,298 genes.

### 3.2 Experimental settings

We performed a leave-one-dataset-out cross validation where 59 datasets were used in the pre-training step and the remaining one dataset was used in the fine-tuning step. This process was repeated 60 times. No label of these 59 datasets was used in the pre-training step. In the fine-tuning step, labels of the remaining one dataset were cross-validated to evaluate the performance. We selected 200 as our encoder output dimension, and each layer in the classifier uses an output dimension that is half of the input vector. When creating pseudo-labels on the encoder output, we trained the encoder for 50 epochs and generated new pseudo-labels once (an iteration), largely due to the time consuming clustering process. We trained the model for 10 iterations, which took us 3 days on a Nvidia RTX 2080 GPU. We chose 1e-4 as the learning rate for Adam optimizer during the pre-training step, and used 30 as the batch size on each dataset. As for the fine-tuning step, we limited the number of data points in the training data to be *min*(1000, 0.6 × *d*), where *d* was the number of data points. This limit allowed a fair comparison among datasets, considering these datasets varied a lot in quantity. In this step we set the learning rate to be 0.002, and deployed an early stop mechanism based on accuracy on the validation set. We used accuracy, Cohen’s Kappa[38], AUPRC and AUROC as metrics to evaluate model classification performance. As for the multi-class case, we regarded each label as a binary-classification problem and calculated AUPRC (AUROC) on this label, and the final result on one dataset was created by taking average AUPRC (AUROC) on all the labels. The results of cell type classification using these metrics are shown below in section 3.3.

In section 3.4, we used K-means to cluster cell representations generated by scPretrain. We used Rand index[39] to show the clustering accuracy, by calculating the consistency between real labels and cluster labels. In addition, we used UMAP[40] to reduce the dimensionality of 1000 data points (our setting of training data number in the fine-tuning step) and then applied it on the whole test dataset. Likewise, we applied K-means to the vectors obtained by UMAP and then calculated its Rand index, which served as the comparison method. In section 3.5, we investigate cell type classification using conventional classifiers. We first used PCA to reduce the dimensionality using 1000 data points of all datasets in order to give a fair comparison. Conventional classifiers were also trained on these data points. We then applied them on the rest of the data to test their performance, which served as the comparison method. In section 3.6, to analyze the performance of scPretrain, we randomly selected training data from the whole fine-tuning dataset and calculated AUPRC of applying neural networks to this selection. This selection and fine-tuning were repeated multiple times to calculate the variance of AUROC. This variance was then used to measure the stability of one dataset.

### 3.3 Cell type classification

We first sought to investigate whether scPretrain can improve cell type classification performance. By evaluating on 60 datasets, we observed significant improvement against the comparison approach that used the same classification method in the fine-tuning step but did not perform pre-training(**Fig. 2**). Overall, our method had significantly better performance on 39, 38, 47, 55 out of 60 datasets in terms of accuracy, Cohen’s Kappa, AUPRC and AUROC respectively. For example, on the Quake Smart-seq2 Spleen dataset, scPretrain achieved 0.93 AUPRC and 0.99 AUROC, while the comparison approach could only get 0.65 AUPRC and 0.72 AUROC. On the Quake Smart-seq2 Heart dataset, scPretrain obtained 0.97 accuracy and 0.95 Kappa coefficient, which were much better than the 0.86 accuracy and 0.79 Cohen’s Kappa of the comparison approach. Averagely, scPretrain got 0.939 accuracy, 0.882 Cohen’s Kappa coefficient, 0.841 AUPRC and 0.975 AUROC on 60 datasets, while the corresponding performance of existing methods was 0.888, 0.747, 0.784 and 0.926. Even though only 39 and 38 datasets achieved significantly better accuracy and Kappa, the performance of scPretrain on the remaining datasets were still comparable to the comparison approach (**Fig. 2a,b**). The substantial improvement of scPretrain on a variety of diverse datasets indicated the effectiveness of performing multi-task self-supervised learning, which learnt a high-quality feature extraction encoder and reduced batch effects across datasets.

**Fig. 2.**
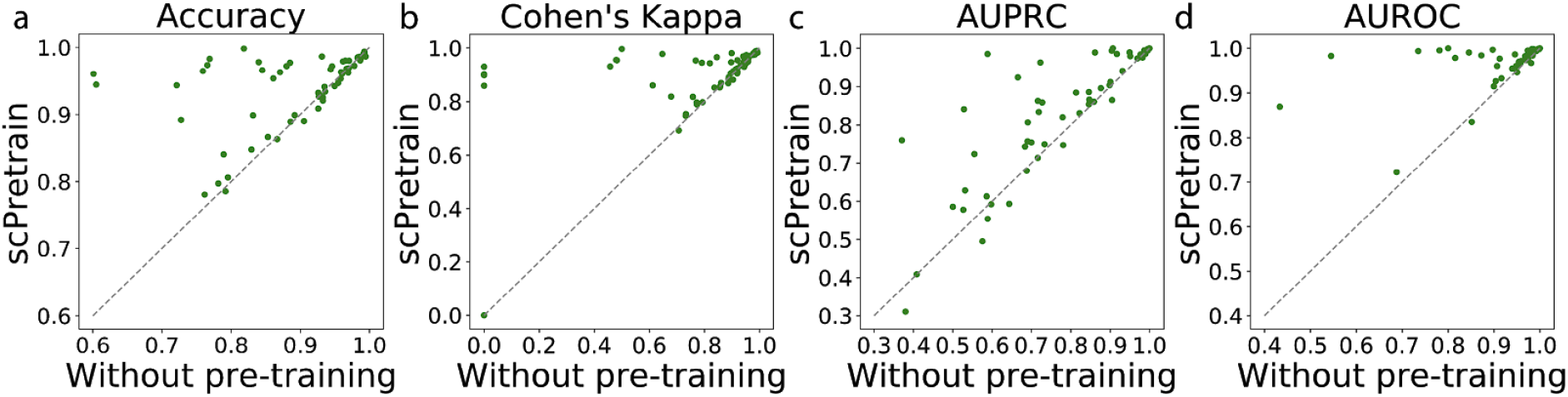
Performance of scPretrain on 60 datasets. a-d, Scatter plots showing the comparison of scPretrain performance and the comparison approach on 60 datasets using accuracy (a), Cohen’s Kappa (b), AUPRC (c) and AUROC (d). Each dot in the plots represents one dataset in the leave-one-dataset-out cross validation. x-axis is the performance of the comparison approach, whereas y-axis is the performance of scPretrain.

### 3.4 Cell clustering using pre-trained cell representation

We studied whether the cell representations obtained using the feature extraction encoder can be used to cluster cells. To this end, we clustered cells using the representations generated from scPretrain without fine-tuning. We found that scPretrain generated better features for cell clustering in comparison to the conventional dimensionality reduction method. scPretrain achieved 0.18 Rand index, which was higher than the 0.15 Rand index of the comparison approach (p-value<0.05). By further projecting cell representations into 2-D space using UMAP, we found that our pre-trained representations tended to form better clusters in agreement to their real cell types (**Fig. 3**). The improved clustering and visualization performance obtained in the pre-training step further raised our confidence about the quality of cell representations created by scPretrain, which was previously verified by the improvement on cell type classification.

**Fig. 3.**
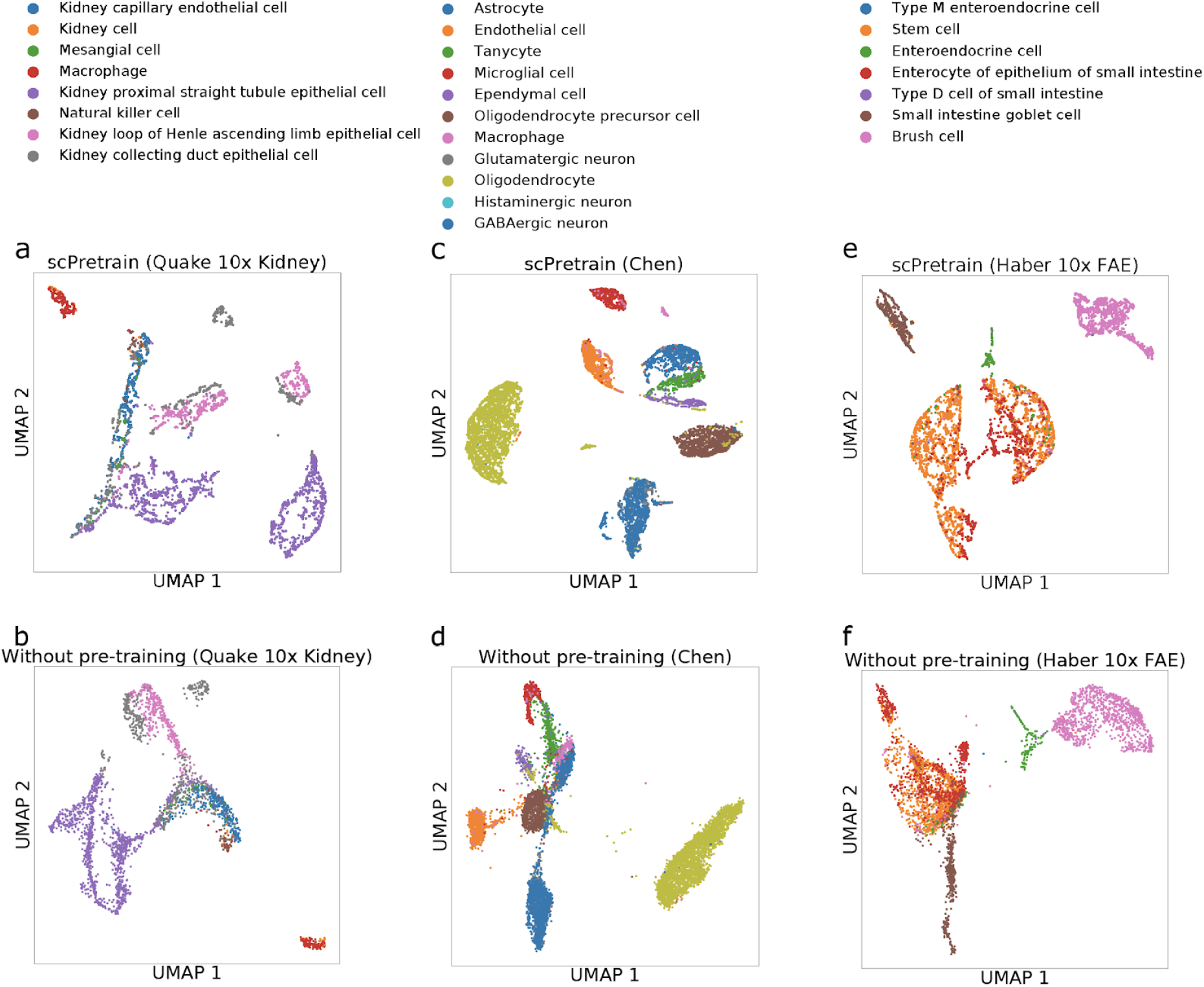
Comparison of cell clustering using scPretrain. a,b,c,d,e,f, Scatter plots showing UMAP visualization of scPretrain (a,c,e) and the comparison approach (b,d,f) on Quake 10x Kidney (a,b), Chen (c,d), Haber 10x FAE (e,f).

### 3.5 Cell type classification using conventional classifiers

We examined the quality of cell representations obtained by scPretrain by using them as features to train other conventional classifiers, including support vector machine (SVM), random forest, and logistic regression. These methods were able to provide more interpretable results and thus revealed marker genes. We found that embeddings using scPretrain again outperformed the comparison approach using all three classifiers on both AUROC and AUPRC (**Fig. 4**). For example, when random forest was used as the classifier, scPretrain outperformed the comparison approach on 95% of all datasets in terms of AUROC and 87% of all datasets in terms of AUPRC (**Fig. 4a,b)**. Likewise, scPretrain outperformed the comparison approach on 85% of all datasets in terms of AUROC and 77% of all datasets in terms of AUPRC using SVM (**Fig. 4c,d)**. Although our pre-training model was mainly devised to provide the neural networks with a stable initialization, it also created robust cell representations that benefited other classifiers as well.

**Fig. 4.**
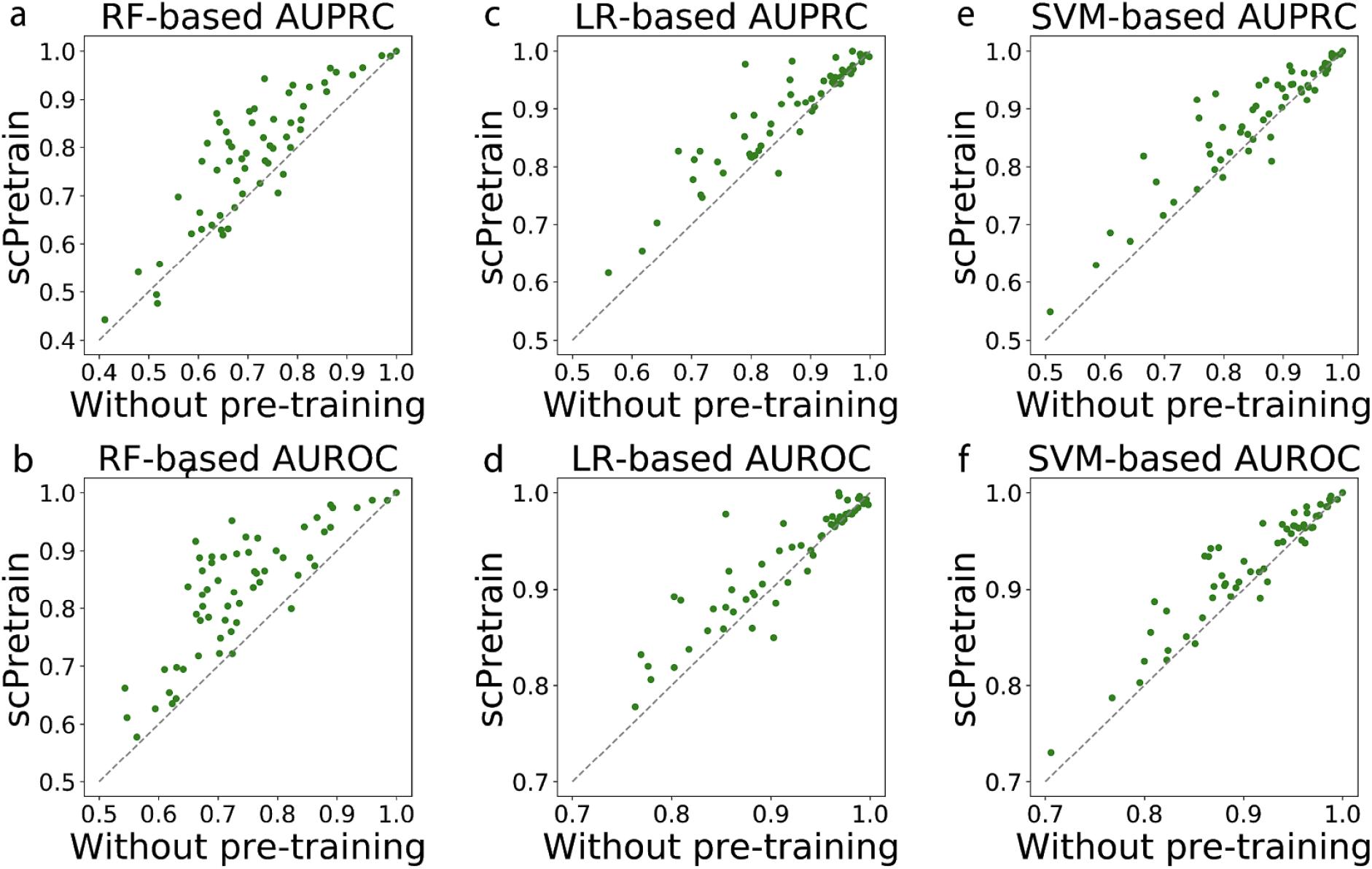
Comparison of the performance using conventional classifiers. a,b,c,d,e,f, Scatter plots showing the performance of using scPretrain generated cell representations as features for random forest (a,b), logistic regression (c,d), and support vector machine (e,f) in terms of AUPRC (a,c,e) and AUROC (b,d,f). As for each plot, each dot represents one dataset. x-axis shows the performance of without pre-training, whereas y-axis shows the performance of scPretrain.

### 3.6 Analysis of scPretrain performance

scPretrain tended to perform well on datasets that standard neural networks were unstable to be trained on. We tested the variance of AUPRC of each dataset using a standard neural network without pre-training and the AUPRC improvement of scPretrain against this standard neural network. We found that these two statistics were highly correlated, with a Pearson correlation coefficient (0.301) (p-value < 0.015). We further observed that the AUPRC improvement was also related to the clustering score improvement in terms of Rand index improvement (Wilcoxon signed-rank test p-value <0.017). Pre-training can be regarded as a regularization strategy by providing the initial parameters[41]. Therefore, the performance of scPretain was more prominent on the dataset that conventional approaches had insatiable performance by injecting strong regularization through pre-training.

## 4 CONCLUSION

In this paper, we have introduced scPretrain, a novel multi-task self-supervised learning method for cell type classification. scPretrain uses pseudo-labels to guide pre-training of a feature extraction encoder using unannotated gene expression profiles. This feature extraction encoder is further fine-tuned using a small number of annotated samples, thus accurately embedding new cells into a low-dimensional space. scPretrain proposes a multi-task learning framework to alleviate batch effects from diverse datasets. We evaluated scPretrain on 60 scRNA-seq datasets and obtained substantial improvement on cell type classification and cell clustering. Also, the robust representations generated by scPretrain have shown to benefit other off-the-shelf classifiers.

In the future, we hope to include more pre-training datasets to further provide users with more robust cell representations using our framework. Pre-training methods in computer vision and natural language processing have been applied to more than 10 millions images and documents, and the performance keeps increasing as the dataset becomes larger. Currently scPretrain’s improvement against the comparison approach decreases when the fine-tuning dataset includes more than 10,000 annotated cells. So by using larger pre-training datasets we hope this framework will be able to further enhance its classification ability on larger fine-tuning datasets. On the other hand, this framework is not confined to single cell expression data, but can be used on other single cell omics data, such as mRNA-DNA methylation, mRNA-chromatin accessibility, and mRNA-protein[28].

## References

1. Klein AM, Mazutis L, Akartuna I, Tallapragada N, Veres A, Li V, et al. Droplet Barcoding for Single-Cell Transcriptomics Applied to Embryonic Stem Cells. Cell. 2015;161: 1187–1201.

2. Guo G, Huss M, Tong GQ, Wang C, Li Sun L, Clarke ND, et al. Resolution of cell fate decisions revealed by single-cell gene expression analysis from zygote to blastocyst. Dev Cell. 2010;18: 675–685.

3. Tabula Muris Consortium, Overall coordination, Logistical coordination, Organ collection and processing, Library preparation and sequencing, Computational data analysis, et al. Single-cell transcriptomics of 20 mouse organs creates a Tabula Muris. Nature. 2018;562: 367–372.

4. Han X, Wang R, Zhou Y, Fei L, Sun H, Lai S, et al. Mapping the Mouse Cell Atlas by Microwell-Seq. Cell. 2018;173: 1307.

5. Tang F, Barbacioru C, Wang Y, Nordman E, Lee C, Xu N, et al. mRNA-Seq whole-transcriptome analysis of a single cell. Nat Methods. 2009;6: 377–382.

6. Grün D, Muraro MJ, Boisset J-C, Wiebrands K, Lyubimova A, Dharmadhikari G, et al. De Novo Prediction of Stem Cell Identity using Single-Cell Transcriptome Data. Cell Stem Cell. 2016;19: 266–277.

7. Muraro MJ, Dharmadhikari G, Grün D, Groen N, Dielen T, Jansen E, et al. A Single-Cell Transcriptome Atlas of the Human Pancreas. Cell Syst. 2016;3: 385–394.e3.

8. Baron M, Veres A, Wolock SL, Faust AL, Gaujoux R, Vetere A, et al. A Single-Cell Transcriptomic Map of the Human and Mouse Pancreas Reveals Inter- and Intra-cell Population Structure. Cell Syst. 2016;3: 346–360.e4.

9. Zheng GXY, Terry JM, Belgrader P, Ryvkin P, Bent ZW, Wilson R, et al. Massively parallel digital transcriptional profiling of single cells. Nat Commun. 2017;8: 14049.

10. Davie K, Janssens J, Koldere D, De Waegeneer M, Pech U, Kreft Ł, et al. A Single-Cell Transcriptome Atlas of the Aging Drosophila Brain. Cell. 2018;174: 982–998.e20.

11. Tabula Muris Consortium. A single-cell transcriptomic atlas characterizes ageing tissues in the mouse. Nature. 2020;583: 590–595.

12. Tan Y, Cahan P. SingleCellNet: A Computational Tool to Classify Single Cell RNA-Seq Data Across Platforms and Across Species. Cell Syst. 2019;9: 207–213.e2.

13. Pliner HA, Shendure J, Trapnell C. Supervised classification enables rapid annotation of cell atlases. Nat Methods. 2019;16: 983–986.

14. Ma F, Pellegrini M. ACTINN: automated identification of cell types in single cell RNA sequencing. Bioinformatics. 2020;36: 533–538.

15. Hou R, Denisenko E, Forrest ARR. scMatch: a single-cell gene expression profile annotation tool using reference datasets. Bioinformatics. 2019;35: 4688–4695.

16. Abdelaal T, Michielsen L, Cats D, Hoogduin D, Mei H, Reinders MJT, et al. A comparison of automatic cell identification methods for single-cell RNA sequencing data. Genome Biol. 2019;20: 194.

17. Lopez R, Regier J, Cole MB, Jordan MI, Yosef N. Deep generative modeling for single-cell transcriptomics. Nat Methods. 2018;15: 1053–1058.

18. Zhang AW, O’Flanagan C, Chavez EA, Lim JLP. Probabilistic cell-type assignment of single-cell RNA-seq for tumor microenvironment profiling. Nature. 2019. Available: https://www.nature.com/articles/s41592-019-0529-1?elqTrackId=12c8cef68e0741ef8422778b61588aec

19. Hu J, Li X, Hu G, Lyu Y, Susztak K, Li M. Iterative transfer learning with neural network for clustering and cell type classification in single-cell RNA-seq analysis. Nature Machine Intelligence. 2020;2: 607–618.

20. Wagner F, Yanai I. Moana: A robust and scalable cell type classification framework for single-cell RNA-Seq data. doi:10.1101/456129

21. Tran HTN, Ang KS, Chevrier M, Zhang X, Lee NYS, Goh M, et al. A benchmark of batch-effect correction methods for single-cell RNA sequencing data. Genome Biol. 2020;21: 12.

22. Haghverdi L, Lun ATL, Morgan MD, Marioni JC. Batch effects in single-cell RNA-sequencing data are corrected by matching mutual nearest neighbors. Nat Biotechnol. 2018;36: 421–427.

23. Shaham U, Stanton KP, Zhao J, Li H, Raddassi K, Montgomery R, et al. Removal of batch effects using distribution-matching residual networks. Bioinformatics. 2017;33: 2539–2546.

24. Li X, Wang K, Lyu Y, Pan H, Zhang J, Stambolian D, et al. Deep learning enables accurate clustering with batch effect removal in single-cell RNA-seq analysis. Nat Commun. 2020;11: 2338.

25. Devlin J, Chang M-W, Lee K, Toutanova K. BERT: Pre-training of Deep Bidirectional Transformers for Language Understanding. arXiv [cs.CL]. 2018. Available: http://arxiv.org/abs/1810.04805

26. Fernando B, Bilen H, Gavves E, Gould S. Self-Supervised Video Representation Learning With Odd-One-Out Networks. arXiv [cs.CV]. 2016. Available: http://arxiv.org/abs/1611.06646

27. Caron M, Bojanowski P, Joulin A, Douze M. Deep Clustering for Unsupervised Learning of Visual Features. arXiv [cs.CV]. 2018. Available: http://arxiv.org/abs/1807.05520

28. Zhang Z, Luo D, Zhong X, Choi JH, Ma Y, Wang S, et al. SCINA: A Semi-Supervised Subtyping Algorithm of Single Cells and Bulk Samples. Genes. 2019;10. doi:10.3390/genes10070531

29. Kimmel JC, Kelley DR. scNym: Semi-supervised adversarial neural networks for single cell classification. Cold Spring Harbor Laboratory. 2020. p. 2020.06.04.132324. doi:10.1101/2020.06.04.132324

30. Dong X, Chowdhury S, Victor U, Li X, Qian L. Cell Type Identification from Single-Cell Transcriptomic Data via Semi-supervised Learning. arXiv [q-bio.GN]. 2020. Available: http://arxiv.org/abs/2005.03994

31. Chen L, He Q, Zhai Y, Deng M. Single-cell RNA-seq data semi-supervised clustering and annotation via structural regularized domain adaptation. Bioinformatics. 2020. doi:10.1093/bioinformatics/btaa908

32. Xu C, Lopez R, Mehlman E, Regier J, Jordan MI, Yosef N. Probabilistic Harmonization and Annotation of Single-cell Transcriptomics Data with Deep Generative Models. doi:10.1101/532895

33. Hu W, Liu B, Gomes J, Zitnik M, Liang P, Pande V, et al. Strategies for Pre-training Graph Neural Networks. arXiv [cs.LG]. 2019. Available: http://arxiv.org/abs/1905.12265

34. Chen M, Radford A, Child R, Wu J, Jun H, Dhariwal P, et al. Generative Pretraining from Pixels. Proceedings of the 37th International Conference on Machine Learning. 2020. Available: https://static.aminer.cn/storage/pdf/icml/20/6022-Paper.pdf

35. Lin Y, Dong X, Zheng L, Yan Y, Yang Y. A bottom-up clustering approach to unsupervised person re-identification. Proc Conf AAAI Artif Intell. 2019. Available: https://www.aaai.org/ojs/index.php/AAAI/article/view/4898

36. Kim T, Chen IR, Lin Y, Wang AY-Y, Yang JYH, Yang P. Impact of similarity metrics on single-cell RNA-seq data clustering. Brief Bioinform. 2019;20: 2316–2326.

37. Cao Z-J, Wei L, Lu S, Yang D-C, Gao G. Searching large-scale scRNA-seq databases via unbiased cell embedding with Cell BLAST. Nat Commun. 2020;11: 3458.

38. Cohen J. A Coefficient of Agreement for Nominal Scales. Educ Psychol Meas. 1960;20: 37–46.

39. Rand WM. Objective Criteria for the Evaluation of Clustering Methods. J Am Stat Assoc. 1971;66: 846–850.

40. McInnes L, Healy J, Melville J. UMAP: Uniform Manifold Approximation and Projection for Dimension Reduction. arXiv [stat.ML]. 2018. Available: http://arxiv.org/abs/1802.03426

41. Erhan D, Courville A, Bengio Y, Vincent P. Why Does Unsupervised Pre-training Help Deep Learning? In: Teh YW, Titterington M, editors. Chia Laguna Resort, Sardinia, Italy: JMLR Workshop and Conference Proceedings; 2010. pp. 201–208.

